# Two-Color Spatial Cumulant Analysis Detects Heteromeric Interactions between Membrane Proteins

**DOI:** 10.1101/613927

**Authors:** D.J. Foust, A.G. Godin, A. Ustione, P.W. Wiseman, D.W. Piston

## Abstract

Fluorescence fluctuation spectroscopy (FFS) can be used to measure the aggregation of fluorescently labeled molecules and is typically performed using time series data. Spatial intensity distribution analysis (SpIDA) and fluorescence moment image analysis (FMIA) are established tools for measuring molecular brightnesses from single-color images collected with laser scanning microscopes. We have extended these tools for analysis of two-color images to resolve heteromeric interactions between molecules labeled with spectrally distinct chromophores. We call these new methods two-color SpIDA (2c-SpIDA) and two-color spatial cumulant analysis (2c-SpCA). To implement these techniques on a hyperspectral imaging system, we developed a spectral shift filtering (SSF) technique to remove artifacts due to intrinsic crosstalk between detector bins. We determined that 2c-SpCA provides better resolution from samples containing multiple fluorescent species, hence this technique was carried forward to study images of living cells. We used fluorescent heterodimers labeled with EGFP and mApple to quantify the effects of resonance energy transfer and incomplete maturation of mApple on brightness measurements. We show that 2c-SpCA can detect the interaction between two components of trimeric G-protein complexes. Thus 2c-SpCA presents a robust and computationally expedient means of measuring heteromeric interactions in cellular environments.

**Statement of Significance:** Fluorescence fluctuation spectroscopy (FFS) techniques determine biophysical parameters from samples containing fluorescently labeled biomolecules by considering the statistical nature of fluorescent signals measured with photodetectors. The present study introduces two-color spatial cumulant analysis (2c-SpCA) to the canon of FFS techniques. 2c-SpCA analyzes pixel-value data of two-color images collected with laser scanning fluorescence microscopes. We show that 2c-SpCA can determine several biophysical parameters in living cells including Forster resonance energy transfer efficiency, the dark state fraction of fluorescent proteins, and heteromerization between distinctly labeled proteins. In comparison to existing techniques, 2c-SpCA requires very few image frames for analysis, minimal computations, and can be applied to images of fixed tissue samples.

## Introduction

Fluorescence fluctuation spectroscopy (FFS) is a set of statistical biophysical techniques commonly used to determine the concentration, transport coefficients, and molecular brightness of fluorescently labeled biomolecules (1). The molecular brightness is of special interest because it contains information about the oligomerization of the molecules studied *in vitro* or in cells. For example, in the absence of quenching, a fluorescent homodimer has twice the brightness of the monomer. Similarly, a fluorescent *n*-mer has *n* times the brightness of the monomer.

To measure the aggregation of fluorescent species, Qian and Elson developed a theory to relate the moments of the fluorescence intensity distribution to the particle number per detection volume and the molecular brightness (2). Moment-based techniques have since been re-expressed as closely related factorial cumulants (FCA) which offer a more compact relationship to number and brightness (3, 4). Later, two groups simultaneously developed methods relating functional forms of photon count distributions to the particle number and brightness (5, 6). These are essentially equivalent techniques (7) and are referred to as photon counting histogram analysis (PCH) and fluorescence intensity distribution analysis (FIDA). Histogram and cumulant approaches have been extended for multicolor analysis, providing improved resolution of heteromeric interactions between molecules labeled with spectrally distinct fluorophores (8–10).

Both histogram and cumulant analyses were originally conceived in the paradigm of time series data from a single detection volume at a fixed position in the sample. These approaches have been successfully modified for analysis of spatially sampled data from scanning systems (confocal and two-photon microscopy). Spatial intensity distribution analysis (SpIDA) is the analog of PCH applied to single images (11, 12) and fluorescence moment image analysis (FMIA) is the corollary to moment analysis of point measurements (13). The advantages of spatial sampling include decreased photobleaching effects due to local fluorophore depletion and the ability to probe slow moving (or static) aggregates that would require very long collection times in a fixed-point configuration. Neither of these approaches have been extended to multicolor analysis.

To extend these approaches for multicolor images, we present two-color spatial intensity distribution analysis (2c-SpIDA) and two-color spatial cumulant analysis (2c-SpCA). We implemented these techniques using a two-photon laser scanning microscope equipped with a 32-bin spectral detector. To circumvent the effects of intrinsic crosstalk between the bins, we implemented a spectral shift filter (SSF) inspired by a temporal shifting approach developed for point measurements (14). With the SSF applied, we show data collected for a single fluorescent species in solution (EGFP) is well-described by ideal detector models of intensity distributions and cumulants. We show that this approach can resolve the molecular brightnesses of multiple species in solution (EGFP and mApple). We determined that 2c-SpCA is the more robust of the two approaches in accordance with previous comparisons of 2c-PCH and 2c-FCA (8, 15). Finally, we applied 2c-SpCA to study fluorescent protein heteromers expressed on the plasma membrane of living cells. After image segmentation to identify homogeneous regions of the cell membrane, we show that 2c-SpCA can accurately determine heteromeric interactions between co-expressed fluorescent protein fusion pairs, both in the presence and absence of Förster resonance energy transfer (FRET). We used this approach to measure coupling between subunits of heteromeric G-proteins and between the G-proteins and a G-protein coupled dopamine receptor.

## Materials and Methods

### Instrumentation

Fluorescence imaging was performed with an LSM 880 (Zeiss, Jena, Germany) equipped with a Chameleon Discovery Ti:Sapphire laser (Coherent, Santa Clara, CA) for two-photon excitation. For experiments exciting EGFP and/or mApple, the laser was tuned to 990 nm. Fluorescence was collected with a 40×, 1.2 NA, C-Apochromat water immersion objective lens. Photons were detected with the Quasar 32-bin spectral detector operating in photon counting mode. Emitted light was separated from excitation using MBS 690+ and directed onto the detector by a diffraction grating such that each bin corresponds to an ~8.9 nm band of the electromagnetic spectrum. Experiments were conducted in ‘Lambda’ mode, meaning an image was collected for each bin. Bins were combined into single- or two-color images in post-acquisition steps described below.

### Intrinsic crosstalk measurement

Low intensity light (<10 kHz/bin) from a 488 nm laser was scattered off the surface of a cover glass. To minimize spatial heterogeneity, we scanned the smallest area allowed by our system, 5.31×5.31 μm^2^. This arrangement provided a stable light source with Poissonian emission detection characteristics. Other scanning parameters included 1024×1024 pixels^2^, 0.52 μs/pixel, and ~5 nm/pixel. Crosstalk was quantified by calculating spatial cross-correlation functions, *G*_*CC*_(ξ, φ), for each pair of spectral bins (16):

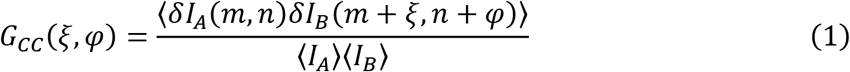

*I*_*A*_ and *I*_*B*_ represent images detected in channels A and B, respectively. δ*I*(*m*, *n*) is a spatial fluctuation defined as *I*(*m*, *n*) − ^〈^*I*(*m*, *n*)^〉^ while *m* and *n* specify image rows and columns, respectively. The lag parameters ξ and φ specify discrete pixel shifts along the scanning and orthogonal axes, respectively. For clarity, we define rows as continuously scanned lines in the raster scanned images and columns as being orthogonal to the scanning direction.

### Imaging-FFS of fluorescent proteins in solution

His_6_-tagged EGFP and mApple were purified using standard immobilized metal affinity chromatography. Briefly, pQE9-EGFP and pQE9-mApple plasmids were transformed into BL21(DE3) *E. coli* (Intact Genomics, St. Louis, MO). Single colonies were used to inoculate 50 mL Luria-Bertani broth supplemented with 100 μg/mL ampicillin and 20 mM glucose. Bacterial pellets were lysed with Bugbuster MasterMix (MilliporeSigma, Darmstadt, Germany) and 1 mM phenylmethylsulfonyl fluoride (PMSF). Fluorescent protein was bound to Ni-NTA Superflow columns (Qiagen, Hilden, Germany) equilibrated with 50 mM NaH_2_PO_4_, 300 mM NaCl, and 10 mM imidazole. Columns were washed twice with 50 mM NaH_2_PO_4_, 300 mM NaCl, and 20 mM imidazole. Purified fluorescent protein was eluted with 50 mM NaH_2_PO_4_, 300 mM NaCl, and 250 mM imidazole.

For imaging, fluorescent proteins were diluted to ~10 nM in PBS (pH 7.4) with 0.01% BSA. Samples were imaged in eight-well chamber slides, #1.5 thickness (ThermoFisher Scientific, Waltham, MA). To avoid adsorption of protein, chamber slides were incubated with 1% BSA for ~30 minutes prior to addition of the sample. Image areas of 10.61×10.61 μm^2^ were scanned with 990 nm excitation (~4 mW at sample). Fluorescence was collected using 23 spectral bins spanning 472-677 nm. Other scanning parameters included: 512×512 pixels^2^, 16.5 μs/pixel, and 21 nm/pixel.

### Plasmids

Construction of pmApple-GNG2 was described previously (17). pEGFP-DRD2 was obtained from Jean-Michel Arrang *via* Addgene (#24099) (18). pEGFP-GNB1 was created by swapping EGFP against mCerulean in pmCerulean-GNB1 (Steven Vogel via Addgene, #27810) (19). pEGFP-HRas was received as a gift from Anne Kenworthy and was originally created by Choy et al (20). pmApple-EGFP-HRas was created by inserting mApple (from pmApple-EGFP described in (17)) at the N-terminus of the EGFP-HRas open reading frame using restriction sites NheI and AgeI. pEGFP(1-233)-mApple(2-236)-HRas was created by inverse PCR of pEGFP-HRas and insertion of mApple(2-236) coding region between EGFP(1-233) and HRas coding regions.

The pCD86(1-19)-EGFP-CD86(20-323)-mApple was created from pCD86-mApple by inserting DNA encoding *LKIPVAT*-EGFP-*SGMAP* (amplified from pEGFP-GNB1) between codons for amino acids 19 and 20 of CD86 in pCD86-mApple. EGFP was not inserted at the absolute N-terminus because the first 19 residues contain a cleavable signal peptide required for normal trafficking to the plasma membrane. The pCD86-mApple had been created by swapping mApple against EGFP in pCD86-EGFP. The pCD86-EGFP had been created by inserting CD86 with a 21 amino acid linker coding region on its C-terminus immediately before EGFP in pEGFP-N1 (Clontech/Takara, Mountain View, CA). The CD86-21aa was amplified from pCD86-mEos2 (Mike Heilemann via Addgene, #98284) (21).

Unless stated otherwise, insertions/swaps were made by amplification of the insert and linearization of the vector by PCR and combination by In-Fusion® enzyme (Takara, Mountain View, CA). Schematics of mature proteins from constructs used in these studies can be seen in Fig. S9.

### Cell culture and live cell imaging

HEK293 cells were grown in 1:1 Dulbecco’s Modified Eagle’s Medium (DMEM)/F-12 Ham with Glutamax+ (ThermoFisher Scientific, Waltham, MA) supplemented with 10% fetal bovine serum (Alkali Scientific, Fort Lauderdale, FL), penicillin, and streptomycin, and incubated at 37 °C with 5% CO_2_. Cells were transfected by electroporation and transferred to glass bottom dishes (30 mm, #1.5 thickness, Cellvis, Mountainview, CA) for imaging. In a typical experiment, ~1.5∙10^6^ cells were electroporated with 1-3 μg of DNA/construct in a 1 mm gap cuvette and divided amongst three dishes. Electroporation was achieved with 8, 150 V pulses lasting 100 μs, separated by 500 ms intervals. Cells were imaged 12-48 hours post-electroporation.

For imaging, growth media was replaced with Kreb’s Ringer Bicarbonate HEPES buffer (KRBH) supplemented with 0.1% BSA and 20 mM glucose. During imaging, cells were kept at 37 °C and 5% CO_2_ using an incubated stage (PeCon GmbH, Erbach, Germany). All images for 2c-SpCA were collected using the same imaging parameters: 3 frames, 512×512 pixels^2^ (21.21×21.21 μm^2^), 132 μs/pixel, 42 nm/pixel, and 990 nm (~1.2 mW at sample).

### Acceptor photobleaching

Pre- and post-bleach acquisitions were made by exciting EGFP and mApple with 990 nm (~1.4 mW at sample). Bleaching was performed using high intensity 561 nm excitation. The imaging field (106.17×106.17 μm^2^) typically contained several cells for analysis. Ten images were collected for each construct tested. ROIs for each cell were drawn by hand. FRET efficiencies for each cell were calculated by: *E* = 1 − *I*_*pre*_/*I*_*post*_. *I*_*pre*_ and *I*_*post*_ are the average measured intensities in the channel (472-517 nm) detecting only donor (EGFP) fluorescence.

### 2c-SpIDA, 2c-SpCA, and HSP analysis

Theory and implementation of 2c-SpIDA, 2c-SpCA, and HSP analysis as well as simulations are described in the Supplementary Material.

### Data analysis

All analyses were performed with scripts written in the Python programming language and are available upon request.

## Results

### Correcting for intrinsic crosstalk in the multianode PMT

Characterization of non-ideal effects in photodetectors is critical to the accuracy of brightness analyses in FFS. For multi-color imaging, hyperspectral approaches have proven efficient, but spectral detectors have additional non-ideal properties that must be accounted for in such measurements. Here we utilized the multianode, 32-bin, Quasar spectral detector on the LSM 880 (Zeiss) operating in photon counting mode. This detector provides excellent flexibility for multicolor imaging experiments, but the placement of multiple anodes on a single monolithic detector can lead to coincident detection of single photons in adjacent spectral bins (22). We noticed non-ideal FFS behavior from this phenomenon, which we refer to as ‘intrinsic crosstalk’ to differentiate it from ‘crosstalk’, which commonly refers to the effects of overlapping emission spectra of fluorescent probes. When adjacent spectral bins are combined post-acquisition, this intrinsic crosstalk has the same effect on brightness analyses as afterpulsing in a single conventional detector as both phenomena result in multiple, apparently simultaneous, detections for a single photon.

The putative mechanism for the generation of intrinsic crosstalk is outlined in Fig. 1A. A single emitted photon strikes the photocathode generating a photoelectron that is subsequently multiplied by a chain of dynodes corresponding to a single spectral bin. During the amplification process a photoelectron may leak into an adjacent bin. The feral photoelectron can undergo its own amplification resulting in the nearly simultaneous detection of the same photon in adjacent anodes.

**FIGURE 1.**
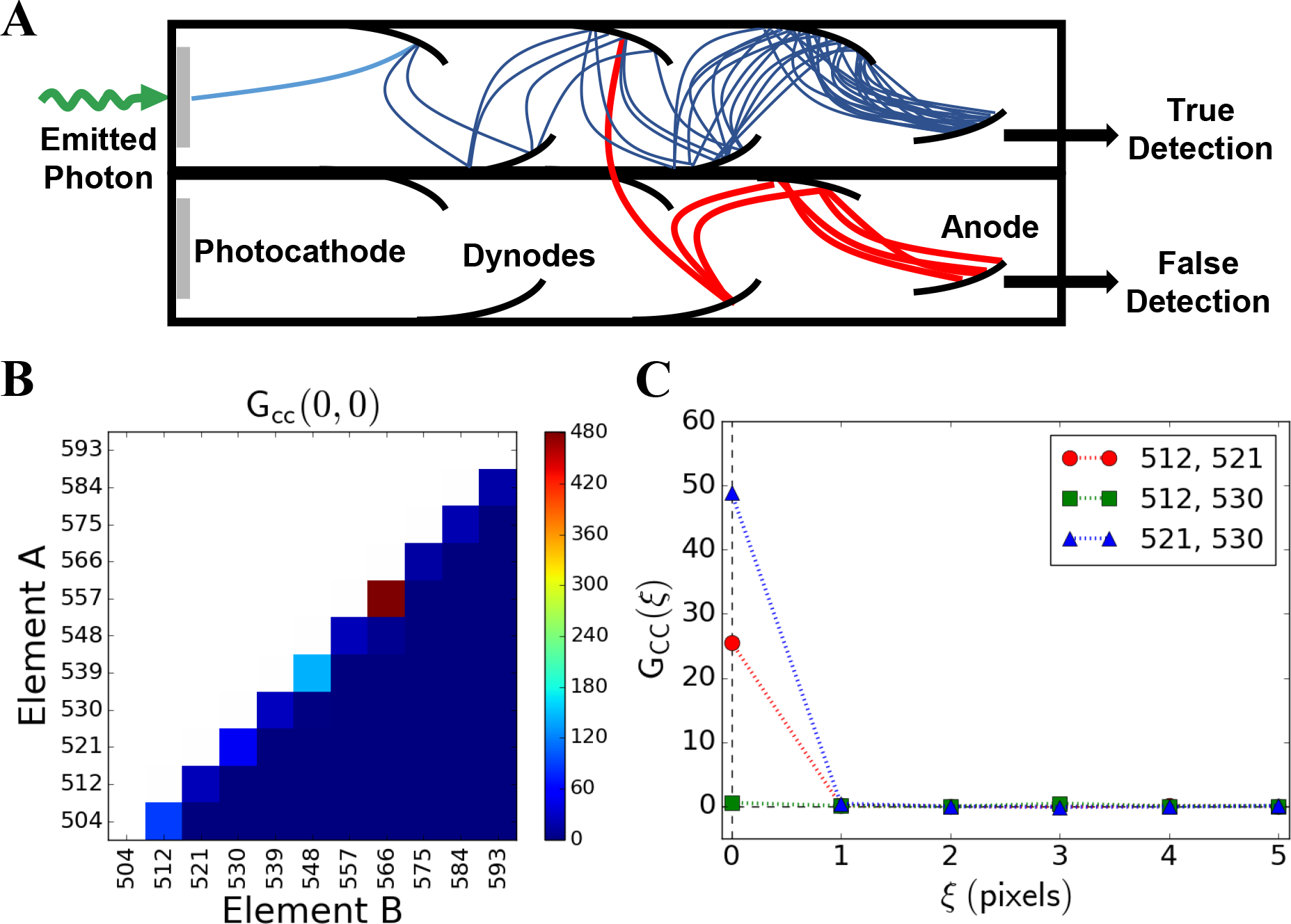
Intrinsic crosstalk between detector elements in a multianode PMT. **(A)** Schematic depicting electronic noise generating a false detection in an adjacent anode. **(B)** Low intensity light was scattered off the surface of a cover glass and detected by channels spanning 504 to 593 nm. The amplitude of the spatial cross-correlation function, G_cc_(0,0), is shown for each pair of detector bins. **(C)** Spatial cross-correlation functions for a single scanning dimension (*G*_*CC*_(ξ, ψ = 0)) for select channel pairs corresponding to emission bands centered at 512, 521, and 530 nm are shown.

To measure intrinsic crosstalk, low-intensity light was scattered off the surface of a glass coverslip while operating in scanning mode. Spatial cross-correlation functions for each pair of detection channels were calculated. In the absence of crosstalk, no correlation is expected between channels measuring a stable light source. Fig. 1B-C show results for 11 channels corresponding to 499-598 nm. We observed large cross-correlation amplitudes between bins that are physically adjacent on the PMT, indicative of intrinsic crosstalk between these channels (Fig. 1B). Conversely, we observed negligible crosstalk between non-adjacent bins. Strong cross-correlation was only present for zero pixel lag (*ξ* = 0, *φ* = 0) between adjacent channels (Fig. 1C), demonstrating that the characteristic timescale of intrinsic crosstalk processes is much shorter than the pixel dwell time (τ_pixel_≈520 ns).

Intrinsic crosstalk poses a significant obstacle to histogram and cumulant analyses of signals measured by our detector. Since its effect on the data are akin to afterpulsing in FFS experiments, we considered several approaches that were developed to address afterpulsing (8, 23–26). These approaches rely on independent measurement of the afterpulsing probability and the modification of models where the afterpulsing probability is treated as a constant. We could not use a similar strategy because an analogous value to the afterpulsing probability for our detector would be dependent on the emission spectra of the sample, not an inherent quality of the detector. For example, a sample with peak emission near 560 nm would likely have a much higher apparent afterpulsing probability than a sample with peak emission near 520 nm due to the very large crosstalk between 557 nm and 566 nm bins (Fig. 1B).

Rather than develop a more complex model to describe intrinsic crosstalk in our system, we opted to introduce a single pixel offset between adjacent channels so that crosstalk could be filtered out in our analyses. Unfiltered images, *I*(*m*, *n*), are constructed from raw images, *i*_*j*_(*x*, *y*), collected in individual bins, *j*, by pixelwise summation of all bins to be included in *I*:

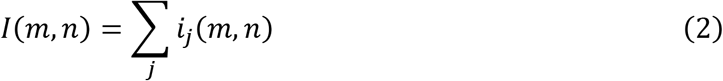

where *m* and *n* specify the row and column of a pixel. Rows are continuously scanned lines during raster scans, whereas columns run orthogonal to the continuously scanned direction. With intrinsic crosstalk, *i*_*j*_(*m*, *n*) and *i*_*j*+1_(*m*, *n*) contain counts corresponding to the same photon, producing an afterpulsing-like effect in *I*(*m*, *n*). Although this phenomenon occurs for a small fraction of detected photons, these contribute to a large deviation from ideal models of photon counting histograms and cumulants. A spectral shift-filtered (SSF) image, *I*_*SSF*_(*m*, *n*), is generated by simply offsetting odd and even indexed channels by one pixel along the scanning axis:

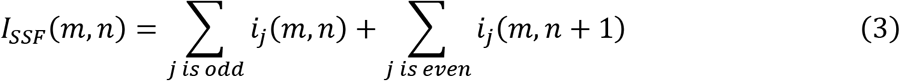

This technique does not introduce significant errors to brightness analyses if two conditions are met: 1) the pixel size is much smaller than the resolution of the optical system, and 2) the pixel dwell time is much smaller than the characteristic diffusion time of the particles being studied. Under these conditions, diffusing particles and the detection volume are approximately stationary over the timescale of one pixel. For immobile particles, only the first condition is relevant. The SSF is illustrated in Fig. S1.

To validate the SSF, we imaged ~10 nM EGFP in PBS and performed several FFS analyses on filtered and unfiltered data. A total of 60 frames of 512×512 pixel^2^ images were collected with 16.5 μs/pixel and 0.021 μm/pixel. EGFP emission was collected in 23 bins spanning 472-677 nm. For single-color analyses, all 23 bins were combined into a single color according to Eq. 2 and Eq. 3 for unfiltered and filtered images, respectively. Likewise, for two-color analyses, the signal was divided as evenly as possible between two channels corresponding to bins covering 472-517 nm (Channel A) and 517-677 nm (Channel B) (Fig. S2). This configuration resulted in an approximately 45/55 split of EGFP fluorescence between channels A and B.

Figs. 2 and 3 show respectively 2c-SpIDA and 2c-SpCA of the 60-frame data set. The efficacy of the SSF can be clearly seen for both analyses. Although it is difficult to observe a large disparity by visual comparison of histograms for unfiltered (Fig. 2A) and SSF (Fig. 2D) data, the difference becomes clear in the normalized residuals (Figs. 2C, 2F) of the fitted histograms (Figs. 2B, 2E) against an ideal detector model. Reduced-χ^2^ for unfiltered and filtered data were 39.9 and 1.17, respectively, clearly indicating that an ideal detector model is appropriate for the SSF data, but not the unshifted data.

**FIGURE 2.**
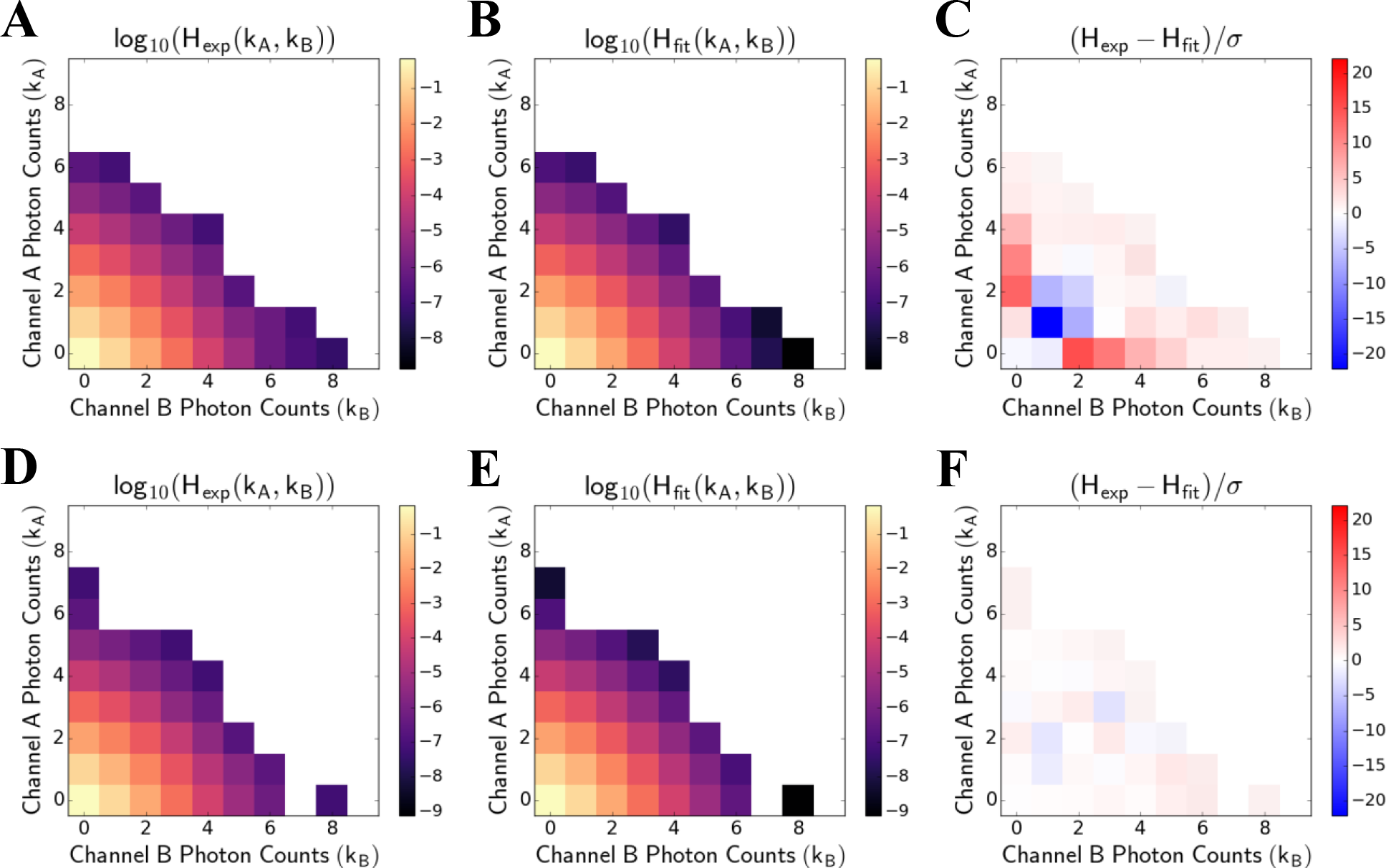
2c-SpIDA for 60 frames of aqueous EGFP with shift filter applied. The top row shows analysis of unshifted data and the bottom row shows analysis with the SSF applied. **(A)** and **(D)** show the experimental bivariate intensity distributions. **(B)** and **(E)** show the fits obtained using a single-component model. **(C)** and **(F)** show normalized residuals of the fits against the experimental distributions. The fitted number density and two-color brightness for unshifted data were 2.13±0.08 and [0.088±0.003, 0.103±0.003], respectively, versus 3.13±0.03 and [0.060±0.001, 0.070±0.001] for SSF data. Reduced-χ^2^ for unshifted and shifted fits were 39.9 and 1.17, respectively. Note: intensity distributions (A-B, D-E) are shown on log scales.

**FIGURE 3.**
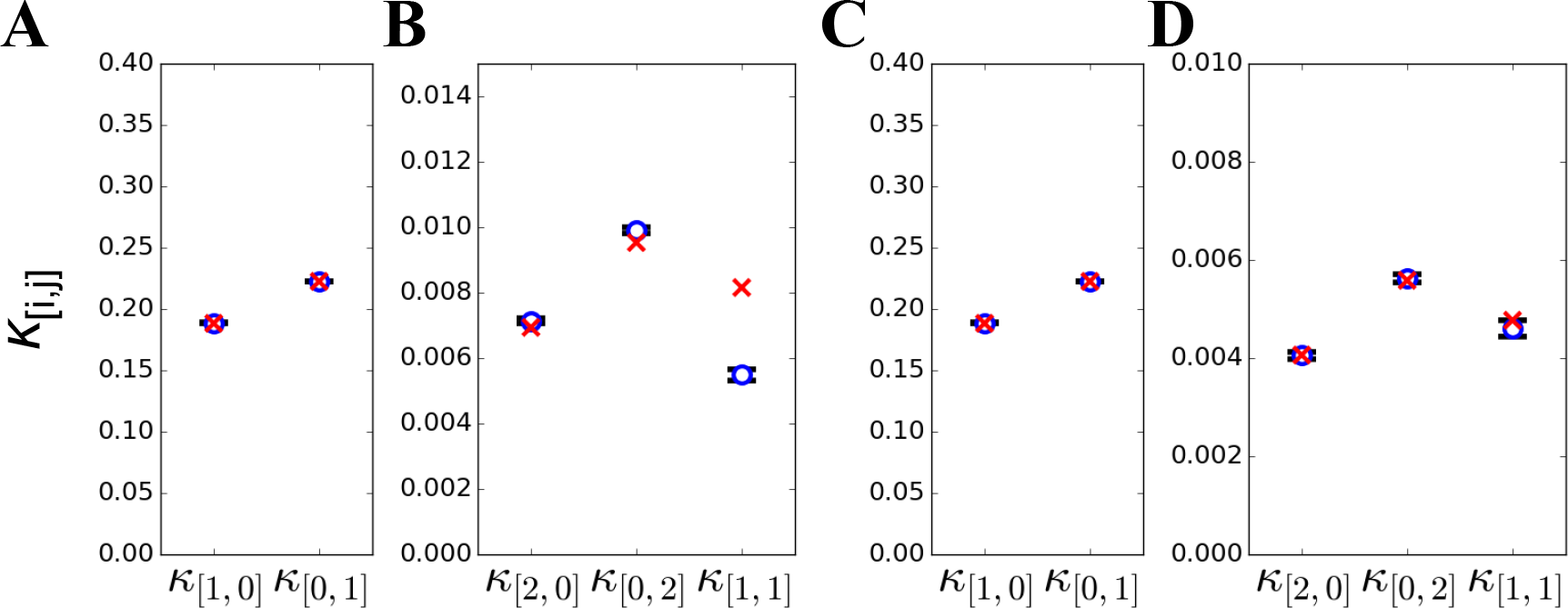
2c-SpCA for 60 frames of ~10 nM EGFP with spectral shift filter applied. The left two panels **(A-B)** show cumulants for unshifted data and the right panels **(C-D)** show cumulants for SSF data. **(A)** and **(C)** show first order factorial cumulants for unfiltered and filtered images, respectively. Likewise, **(B)** and **(D)** show second order factorial cumulants. Experimental factorial cumulants are shown as blue circles and fitted cumulants as red exes. Error bars show estimated uncertainty based on moments-of-moments calculations. Reduced-χ^2^ for unshifted and shifted fits were 127 and 0.58, respectively.

For 2c-SpCA, the SSF has no impact on the first order cumulants (Figs. 3A, 3C), but its effect on the second order factorial cumulants is apparent (Figs. 3B, 3D). Doubly counted photons make a large contribution to the univariate cumulants of individual channels (increase *κ*_[2,0]_, *κ*_[0,2]_) but are weakly correlated between channels (little effect on *κ*_[1,1]_). The single-component, ideal detector model fits the factorial cumulants of the SSF data well but does not fit the unfiltered data. Reduced-χ^2^ values were 127 and 0.58 for unfiltered and SSF fits, respectively, corroborating the suitability of the model for SSF data.

To evaluate the SSF data, we compared different single-color FFS analyses including raster image correlation spectroscopy (RICS), SpIDA, and SpCA. Using RICS, we could determine the molecular brightness from unfiltered data without accounting for intrinsic crosstalk by using the common approach of excluding *G*(0,0) when fitting the spatial autocorrelation function (Fig. S3A). The molecular brightness determined using RICS on unfiltered data closely matched molecular brightnesses determined using 1c-SpIDA (Fig. S3B) or 1c-SpCA on SSF data (Fig. S4). The consistency between RICS analysis of unfiltered data and other analyses of SSF data confirmed that the single pixel offset does not introduce a systematic error for the imaging parameters used. Single-color brightness analyses were also consistent with two-color analyses (Fig. S4).

A comparison of brightnesses determined by single- and two-color methods on shifted and unshifted data is shown in Fig. S4. Analyses where intrinsic crosstalk could be ignored (SSF or RICS) gave a brightness of ~0.13 cpm, whereas crosstalk influenced measurements returned ~0.19 cpm. The apparent increase in in molecular brightness due to intrinsic is accompanied by a proportional decrease in the apparent number density (~3.1 particles/volume with SSF versus ~2.13 unfiltered). The nearly 50% error caused by intrinsic crosstalk was consistent across analyses.

### Resolving fluctuations from EGFP and mApple in solution

We verified that the offset filter in conjunction with 2c-SpIDA or 2c-SpCA could be used to resolve a sample containing multiple fluorescent components by imaging ~10 nM EGFP and mApple individually and together in aqueous solution. Fluorescence emission was divided into two channels to maximize separation of the two chromophores. Channels A and B were constructed from bins spanning 472-562 nm and 579-678 nm, respectively (Fig. S5). Single-component, ideal detector models were applied to SSF data of samples containing a single species and two-component models were applied to the EGFP/mApple mixture. To reduce the number of free parameters in the two-component fit, the brightness of component 2 (mApple) in Channel A was fixed to 0 (*ε*_2,*A*_ = 0) for both 2c-SpIDA and 2c-SpCA. This approximation could be made due to the vanishingly small fluorescence of mApple in Channel A.

Results for 60 frame acquisitions divided into 6 frame segments analyzed individually are shown in Fig. 4. 2c-SpIDA and 2c-SpCA returned nearly identical results. From SpCA, EGFP alone had an average brightness of [0.118, 0.003] cpm versus [0.117, 0.004] for component one (C1) of the mixture. Similarly, the average molecular brightness of mApple alone was [0.001, 0.050] versus [0 (fixed), 0.053] for component two (C2) of the mixture.

**Figure 4.**
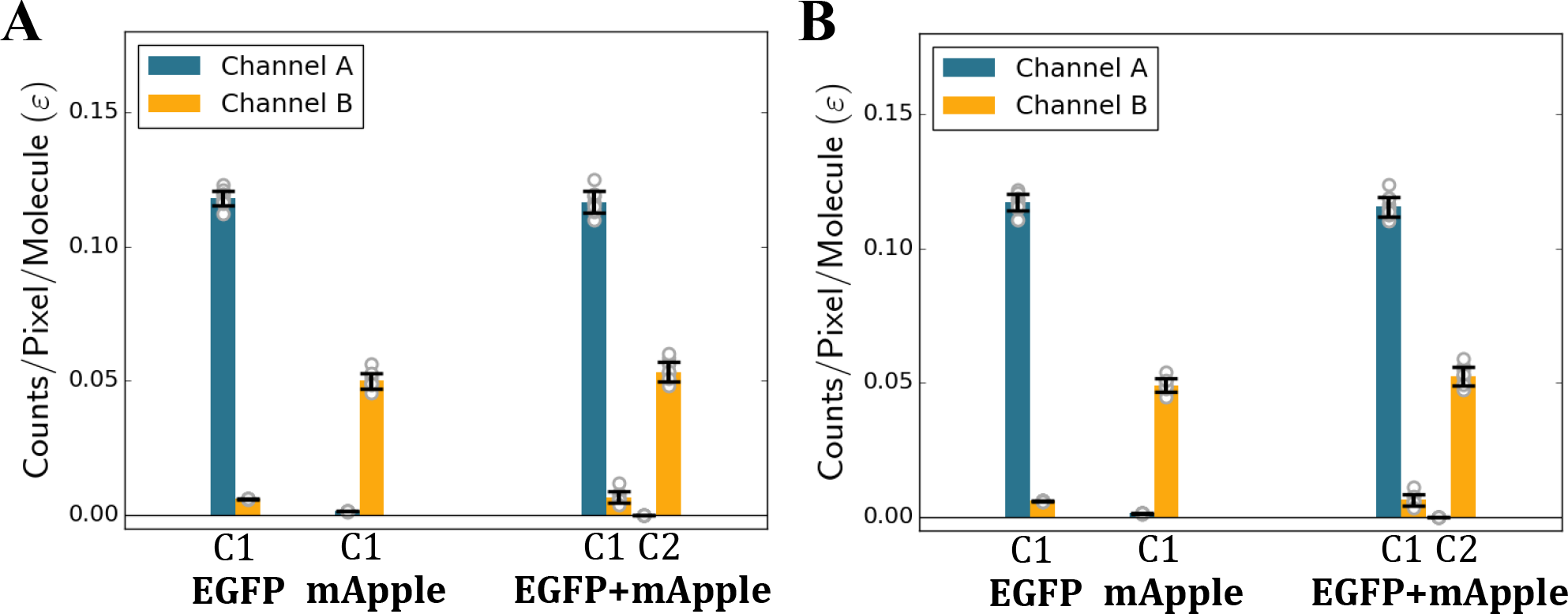
Resolution of aqueous EGFP and mApple mixtures. ~10 nM EGFP and mApple were imaged separately or in a mixture. 60 frame acquisitions were divided into 10 segments of 6 frames. Molecular brightnesses were determined for each segment by 2c-SpCA **(A)** and 2c-SpIDA **(B)**. A single-component model was used to fit EGFP or mApple alone. A two-component model was used to fit the mixture. To reduce the number of free parameters, the brightness of component two (C2) in channel A was fixed to zero (ε_2,A_=0). Channels A and B include fluorescence detected from 472-562 nm and 579-678 nm, respectively.

### 2c-SpIDA versus 2c-SpCA

Although 2c-SpIDA and 2c-SpCA are essentially equivalent methods in solution when large datasets can be obtained without consideration for photobleaching or cellular movements (Fig. 4), we chose to use 2c-SpCA analysis in cells for several reasons. Wu et al (8) showed that the resolution of a two-component mixtures in cells by 2c-FCA is very weakly dependent on the total concentration of the chromophores. In contrast, the resolution of two-component mixtures by 2c-PCH is diminished as concentrations are increased (15). Since 2c-PCH required larger data sets (longer collection times) to achieve the same resolution. These findings suggested that 2c-FCA is the more robust of the two methods.

We recapitulated these results for spatially sampled data by fitting a single-component model to our data of EGFP and mApple in solution. The 60-frame acquisition was iteratively subdivided into smaller segments. The quality of single-component fits was assessed by calculating the reduced-χ^2^. If the data are insufficient to resolve a two-component mixture, we expect to obtain reduced-χ^2^≈1. Since the sample contains both EGFP and mApple, we expect the single-component model to yield a reduced-χ^2^>>1. The results of this analysis are shown in Fig. S6. For large sample sizes (M>10^5^ pixels), both 2c-SpIDA and 2c-SpCA have reduced-χ^2^ much greater than one, indicating that the single-component model is inadequate. For smaller samples, 2c-SpIDA has a reduced-χ^2^ near 1 whereas 2c-SpCA continues to have reduced-χ^2^ that are consistently greater than 1 with as few as ~2∙10^4^ pixels. This indicated that 2c-SpCA is better able to resolve EGFP and mApple with smaller data sets at these concentrations.

Cumulant analysis has the advantage of having a more intuitive visual interpretation than histogram analysis. To highlight this advantage, we directly calculated the second order factorial cumulants and bivariate intensity distributions for an EGFP/mApple mixture with different degrees of heteromerization (Fig. S7). As the degree of heteromerization is increased, there is a clear change in the second order bivariate cumulant (*κ*_[1,1]_) as the degree of heteromerization is increased (Fig. S7A). In contrast, comparison of intensity distributions for different degrees of heteromerization does not provide any easily observable differences (Fig. S7B-D). Bivariate intensity distributions require model fitting to learn useful information about the sample whereas their corresponding cumulants provide directly observable information.

We considered the computation time required for cumulant and histogram analyses. We timed fitting procedures for bivariate histograms and cumulants calculated from the same data. Data were obtained from images of cells expressing EGFP-HRas (see below for details). To fit a single-component model we found 2c-SpIDA took ~1 s versus <10 ms for 2c-SpCA (Fig. S8). On average, 2c-SpCA was ~270× faster than 2c-SpIDA. We expect this difference in computation time to become starker for multicomponent model fitting because 2c-SpIDA requires successive convolution of single-component histograms, whereas cumulants are simply additive for multiple components.

### Region of interest selection for measurements on plasma membrane

In FFS, the fluorescence signal is typically assumed to be stationary. For imaging-FFS, this assumption would require the concentrations and molecular brightnesses of the fluorescent species to be independent of position in space. In practice, rigorously stationary signals are unobtainable from images of single cells, so we settle for finding regions of interest that appear homogeneous. By ‘homogeneous’, we mean lacking obvious visible trends including gradients, puncta, dim spots, etc. For these studies, we sought homogeneous regions from images of plasma membrane-associated proteins in HEK293 cells. To do this, we implemented a segmentation algorithm based on one developed for arbitrary-region RICS (27).

Three frames were collected per acquisition to provide adequate statistics and avoid excessive photobleaching. Prior to segmentation, two-color images were constructed from raw spectral images with the SSF applied (Eq. 3). Channels designated by the same configuration used for the EGFP/mApple resolution study above (Channel A: 472-562 nm, Channel B: 579-678 nm). For segmentation, an initial ROI was drawn to denote a single cell and a spatial averaging filter was applied. We used a disk-shaped kernel with an 11 pixels radius to generate the spatially averaged images. Upper- and lower-pixel value thresholds were chosen for each channel and applied to the spatially averaged images. These thresholds were chosen so that bright spots from accumulation of fluorescent proteins on intracellular organelles would be excluded. The same thresholds were used to generate unique ROIs for all three frames. Finally, the overlap between the thresholded regions of Channels A and B was used to define the final ROI for 2c-SpCA. Fig. 5 summarizes this algorithm for a cell expressing EGFP-CD86-mApple (G-CD86-A, Fig. S9G). This approach was used for the cellular imaging-FFS experiments described below.

**Figure 5.**
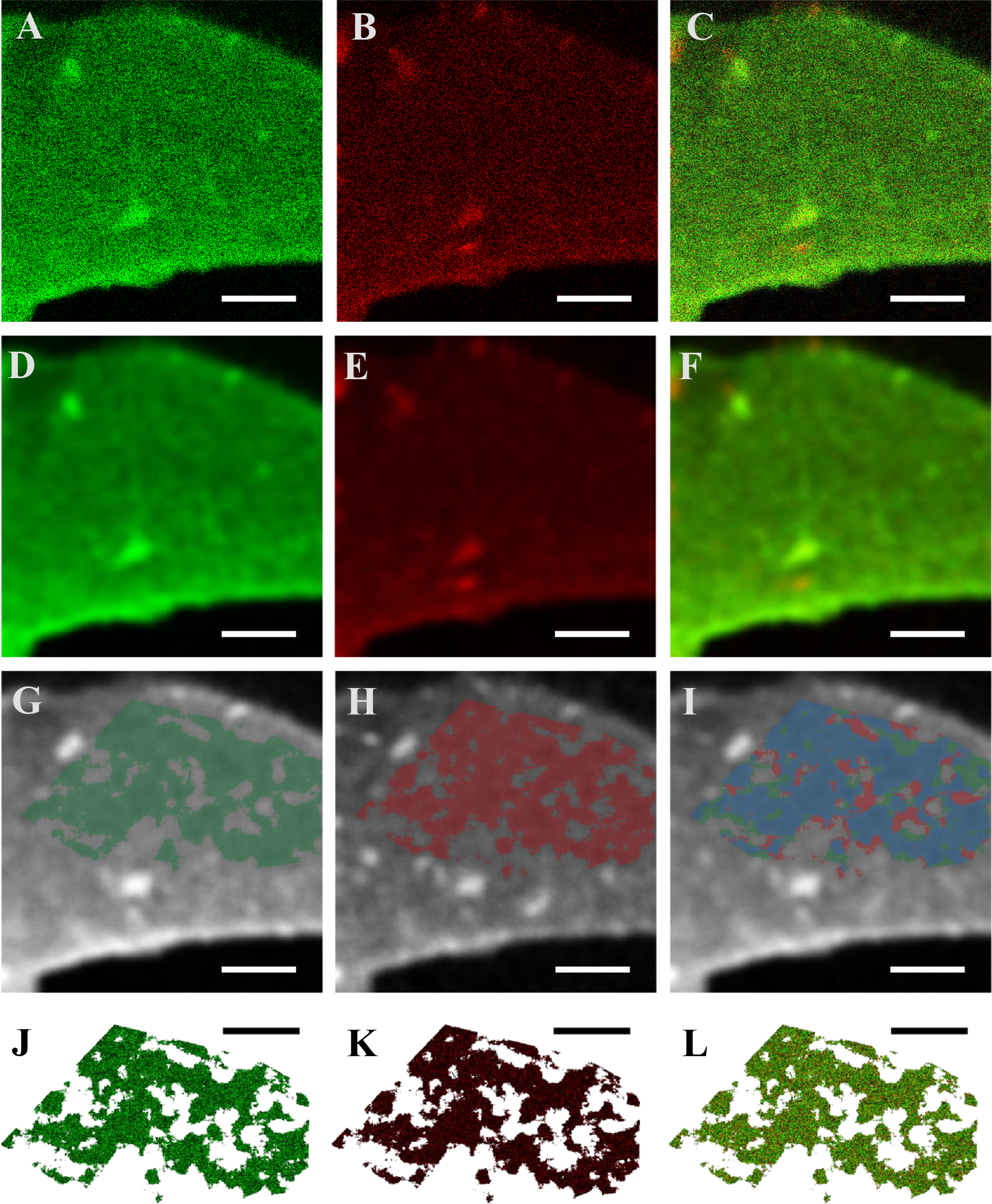
ROI selection for two-color analyses. Images show basal plasma membrane of a HEK293 cell expressing G-CD86-A. The left column corresponds to fluorescence emission from 472-562 nm (channel A) and the middle, 579-678 nm (channel B). The right column shows the channels merged. **A-C** show images after the shift filter was applied. **D-F** show a spatial averaging filter applied to the images using a disk-shaped kernel with an 11 pixel radius. ROIs for each channel are selected by choosing lower and upper bounds for pixel values in the averaged images. The highlighted regions in **G** and **H** show ROIs selected by choosing thresholds for **D** and **E**, respectively. The blue highlighted region in **I** shows the overlap of ROIs selected for the individual channels. Highlighted ROIs in **G-I** are overlaid on grayscale versions of **D-F**. **J-L** show the same data as **A-C** with only the pixels falling within the overlapping ROI (blue region in **I**) shown. Pixel values from **J** and **K** are used in brightness analyses. Although a single frame is shown, typically thresholds were chosen for a three frame sequence simultaneously and pixel values for all three frames were combined for improved statistics. The scale bar is 5 μm for all images.

### 2c-SpCA of EGFP and mApple heterodimers in cells

As our initial test of 2c-SpCA in living cells, we investigated three constitutive heterodimers. Because FRET can be a confounding effect in two-color FFS, we used constructs with different FRET efficiencies tuned by varying the distance between EGFP (G) and mApple (A) chromophores (Fig. S9). A heterodimer with negligible FRET efficiency was constructed by placing EGFP near the N-terminus of CD86 and mApple at the C-terminus (G-CD86-A, Fig. S9G). In this arrangement, EGFP is extracellular and mApple is intracellular in the mature protein. Additional spacing between chromophores is provided by two extracellular, globular domains of CD86.

Low and high FRET efficiency constructs were created by putting EGFP and mApple at the N-terminus of HRas. For the low FRET efficiency construct, mApple was placed at the absolute N-terminus, an 18 amino acid linker separated mApple from EGFP, and 4 amino acid linker separated EGFP from HRas (A-G-HRas, Fig. S9B). For the high FRET efficiency construct, EGFP was placed at the absolute N-terminus, followed by mApple, and HRas (G-A-HRas, Fig. S9C). No linker peptides are present in G-A-HRas, and to further reduce the distance between the EGFP and mApple chromophores the final six residues of EGFP and the first residue of mApple were deleted. These deletions could be made without affecting the photophysics of the chromophores because the C-terminus of EGFP is known to be disordered and the N-terminal methionine of mApple is not essential (28). We verified the relative FRET efficiencies of G-A-HRas, A-G-HRas, and G-CD86-A using acceptor photobleaching (Fig. 6A). G-CD86-A, A-G-HRas, and G-A-HRas had apparent FRET efficiencies of −0.02±0.07, 0.26±0.04, 0.45±0.03, respectively.

**Figure 6.**
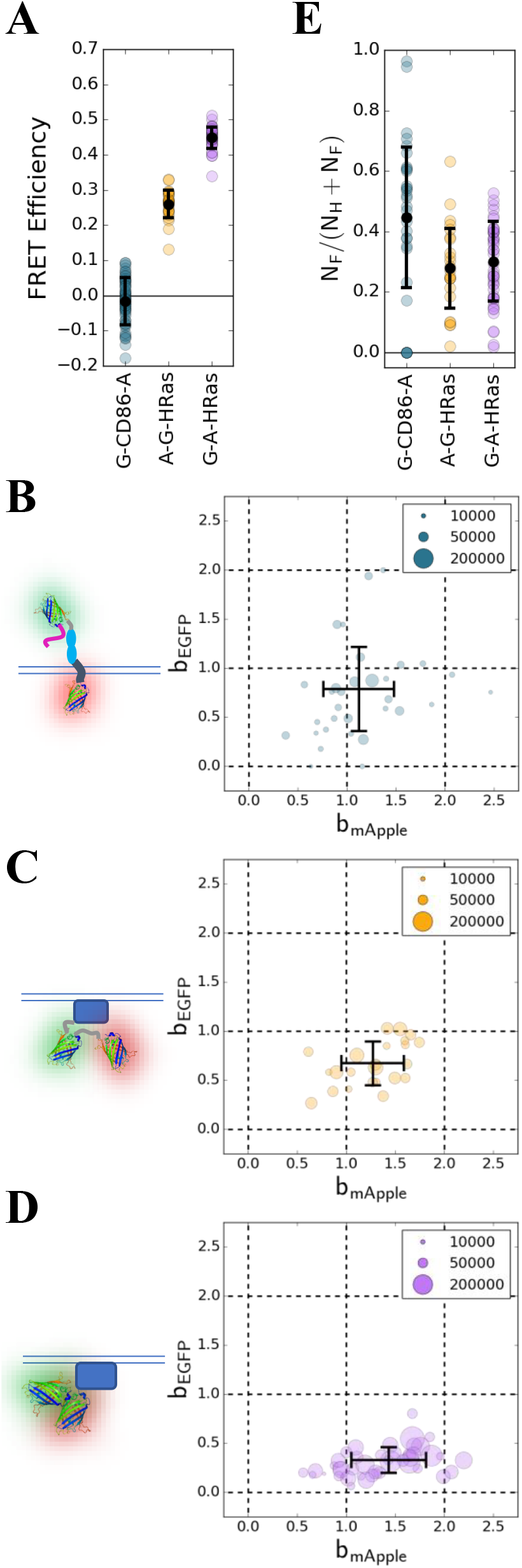
2c-SpCA of constructs constitutively expressing EGFP and mApple. **(A)** The FRET efficiencies of constructs bearing EGFP and mApple determined by acceptor photobleaching. 2-component brightness of the heterospecies determined by 2c-SpCA for: **(C)** G-CD86-A, **(D)** A-G-HRas, and **(E)** G-A-HRas. Normalized brightness plots are accompanied by schematics of the constructs that were imaged. The legends relate the size of the ROI obtained for each cell (in pixels) to the size of the circles in the plot. **(E)** Estimated non-fluorescent fraction of mApple based on 2c-SpCA heterospecies partition analysis. N_F_ and N_H_ are the number densities of the free and heterospecies, respectively for the majority EGFP population. For **(A)**, the error bars denote the mean and one standard deviation. For **(B)-(E)**, the error bars denote the ROI size-weighted (in pixels) mean and standard deviation.

We then performed imaging and 2c-SpCA with these probes. Basal plasma membranes (adjacent to the cover glass) were imaged for three frames. SSF, segmentation, and 2c-SpCA analysis were performed as described above. Although single constructs were transfected into HEK293 cells for these studies, we opted to use a two-component model to account for well-known issues including photobleaching and incomplete maturation.

To study these constructs with a two-component model, we implemented heterospecies partition (HSP) analysis described by Wu et al (29). The HSP analysis we implemented assumes 1:1 binding stoichiometry between molecules so that the sample can be assumed to contain three species (i.e. G, A, G-A). Three species can be reduced to two by defining free and heterospecies. This simplification is necessary to reduce the number of free parameters during the model fitting. The heterospecies is chosen to correspond to the independent fraction of the less abundant chromophore (G or A) in union with the heteromeric species (G-A). The remaining contribution to the factorial cumulants is assigned to the independent fraction of the chromophore that is more abundant (A or G). For example, if the green chromophore is in excess (i.e. [G]+[G-A]>[A]+[G-A]), then the free species corresponds to the independent green chromophore (G) and the heterospecies corresponds to the union of independent red chromophore and the red-green heteromer (A∪G-A). For these experiments, we assumed EGFP chromophores were in excess for two reasons. First, we noticed that mApple was less stable than EGFP when excited with 990 nm, so photobleaching while searching for an appropriate cell and during acquisition primarily reduces the number of detected mApple chromophores. Second, careful inspection of the emission spectra of mApple when excited with 990 nm reveals a small peak at ~510 nm (Fig. S5). The presence of this peak suggests that a population of mApple chromophores exists in a dim, immature state that emits green fluorescence. Brightnesses were normalized to the brightnesses of independently measured monomeric controls. G-HRas and A-GNG2 were used as monomeric controls for EGFP and mApple, respectively.

The two-color normalized brightnesses for the heterospecies of G-CD86-A, A-G-HRas, and G-A-HRas are shown in Fig. 7B-D. In these normalized brightness plots, an ideal heteromer appears at position (1,1). This is the situation where the heterospecies is composed entirely of G-A and energy transfer does not occur. For ideal, non-interacting molecules, the heterospecies appears at (0,1). By design, the heterospecies in these experiments is expected to be almost entirely composed of complete heteromer because independent mApple is unlikely to occur considering the superior maturation and photostability of EGFP. Deviations from the ideal (1,1) position are primarily due to FRET.

**Figure 7.**
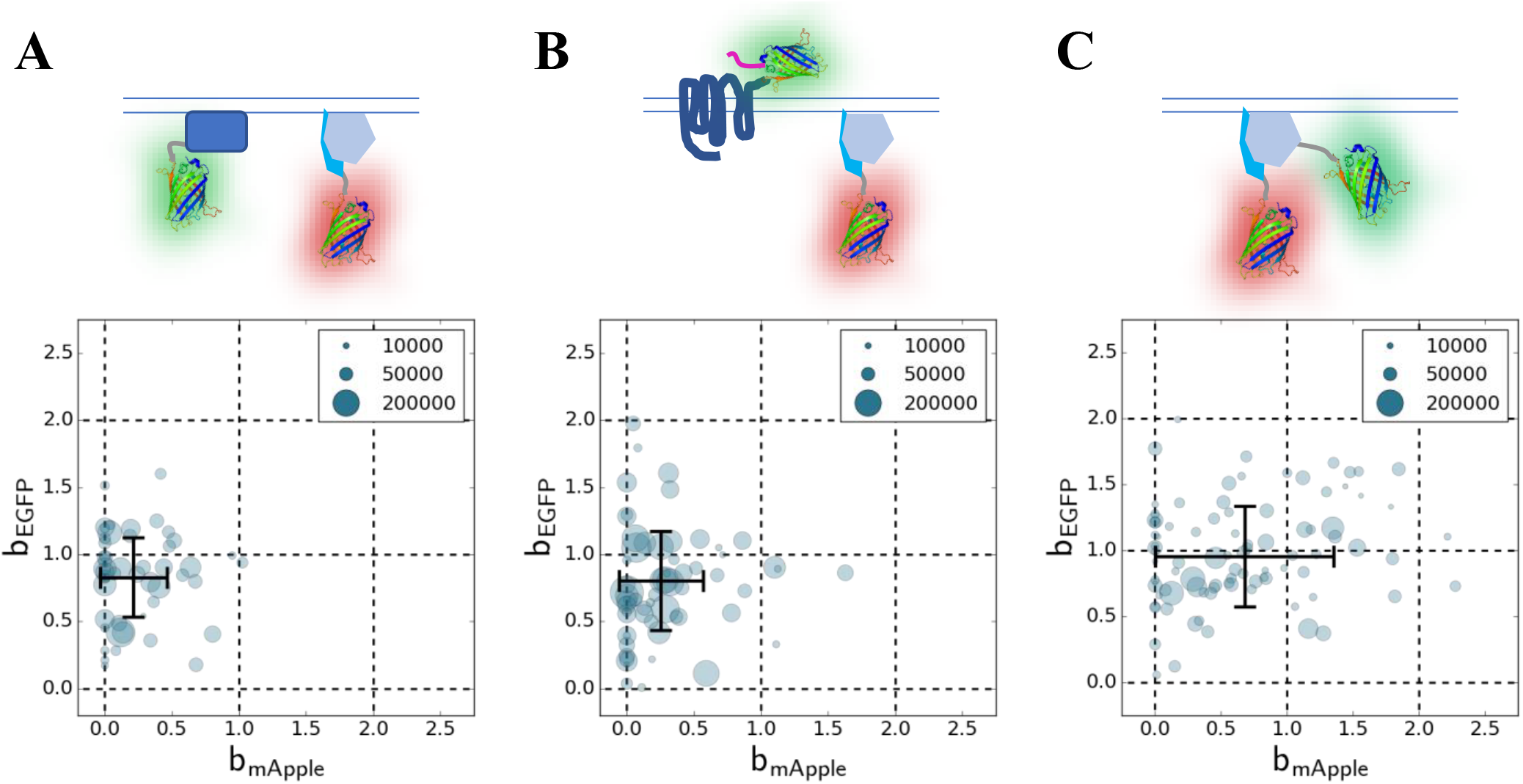
Normalized brightnesses of the heterospecies for cotransfected protein pairs. 2c-SpCA was performed on images of three pairs of fluorescent proteins: **(A)** G-HRas and A-GNG2, **(B)** G-DRD2 and A-GNG2, **(C)** G-GNB1 and A-GNG2. The normalized brightness of the heterospecies from HSP analysis is shown. The key relates the size of the ROIs obtained for each cell in pixels to the size of the circles in the plot. The black cross indicates the ROI size-weighted average of the normalized brightness. Similarly, the error bars indicate the ROI size-weighted standard deviation.

The effect of FRET on the brightness of the heterospecies appears as quenching of EGFP-like brightness, b_EGFP_, and an increase of the mApple-like brightness, b_mApple_. This can be seen for the three constructs we studied (Fig. 6B-D). The higher FRET efficiency construct (G-A-HRas) has a larger down and rightward displacement from (1,1) than the lower FRET efficiency construct (A-G-HRas). We also observe a small displacement for our FRET-free construct (G-CD86-A) on average, however, this likely reflects experimental uncertainty more so than energy transfer that was not detected by acceptor photobleaching. The downward displacement from (1,1) (quenching of EGFP) is a measurement of the FRET efficiency of the heteromer. The average FRET efficiencies by this method for A-G-HRas and G-A-HRas were ~0.33 and ~0.67, respectively. These values are both greater than corresponding measurements made by acceptor photobleaching (Fig. 6A). This discrepancy is explained by the fact that acceptor photobleaching measures sensitized emission from all *potential* donor chromophores including those fused to non-fluorescent (and non-quenching) acceptor molecules. Therefore, acceptor photobleaching tends to underestimate the FRET efficiency when compared to a measurement of a *pure* heteromer population (non-fluorescent acceptors excluded). Significantly, 2c-SpCA offers the ability to accurately measure FRET efficiencies of heteromers in the presence of non-fluorescent acceptors.

We estimated the fraction of mApple chromophores in dim or non-fluorescent states using the number densities of free and heterospecies determined by 2c-SpCA and HSP analysis. Because we assumed the population of independent mApple is small ([A]≈0), we can estimate the fraction of EGFP molecules that do not have an mApple accomplice by: *N*_*F*_/(*N*_*F*_ + *N*_*H*_) where *N*_*F*_ and *N*_*H*_ are the number densities of the free and heterospecies, respectively. This fraction is approximately equal to the fraction of mApple molecules in dark states (Fig. 6E). The two HRas constructs we tested had similar apparent fractions of mApple in dark states (~0.25), whereas G-CD86-A had a greater apparent dark state fraction (~0.45). The larger dark state fraction of G-CD86-A could reflect decreased maturation efficiency of mApple as a C-terminal tag.

### 2c-SpCA of co-expressed of plasma membrane proteins

The most salient application of two-color FFS is the detection of heteromeric interactions between species labeled with spectrally distinct chromophores. We tested the ability of 2c-SpCA to detect interactions between three pairs of plasma membrane proteins: G-HRas/A-GNG2, G-DRD2/A-GNG2, and G-GNB1/A-GNG2. HRas and GNG2 are not known to interact and serve as a negative control. GNG2 and GNB1 form the Gβγ-subunit of the trimeric G-protein complex and form constitutive heterodimers. DRD2 is a G-protein coupled dopamine receptor and is known to transiently interact with trimeric G-protein complexes, including those containing GNG2 and GNB1.

Imaging data of HEK293 cells expressing these protein pairs were fit with a 2-component model using HSP analysis. In contrast to the studies of constitutive heteromers above, HSP analysis was implemented assuming mApple chromophores were in excess. With this configuration, the heterospecies is covers independent EGFP chromophores and heteromeric complexes (G∪G-A) and the free species covers independent mApple (A). In normalized brightness plots (Fig. 7), the heterospecies appears at (0,1) for ideal non-interacting proteins and (1,1) for ideal heteromers. The value of the first coordinate of the heterospecies brightness, *b*_*mApple*_, is a measurement of the degree of heteromerization.

To ensure mApple chromophores were more abundant than EGFP, we checked the ratio between the average signals in Channels A and B. Since EGFP is ~2.4× brighter than mApple (Fig. 4), we limited our analysis to cells where the average signal in Channel B was >1× the average signal in Channel A. This condition made it so that concentration of mApple chromophores would be more than twice the concentration of EGFP chromophores in the absence of FRET. The effect of FRET would be to boost the apparent concentration of mApple and depress the apparent concentration of EGFP. In the presence of moderate FRET, our condition is robust enough guarantee excess mApple. For the fluorescent fusion protein pairs studied, large FRET efficiencies were very unlikely.

For G-HRas/A-GNG2, the normalized brightnesses of the heterospecies determined by 2c-SpCA tended to cluster near (0,1), but we did observe a significant amount of background so that the ROI-size weighted mean deviated from (0,1) (Fig. 7A). For G-DRD2/A-GNG2, fewer normalized brightness measurements were clustered near (0,1) in comparison to G-HRas/A-GNG2, but on average these two pairs were not clearly differentiable by visual inspection (Fig. 7B). Conversely, normalized brightnesses from cells expressing G-GNB1/A-GNG2 show clear heteromerization (Fig. 7C).

We compared our results with simulations of EGFP and mApple with different degrees of heteromerization over a range of ROI sizes, as detailed in the Supplementary Information (Fig. S10). Qualitatively, our experimental results exhibit greater variance than would be expected for similarly sized simulated ROIs. This observation is not unexpected considering that simulations do not capture several sources of variance. For example, our ROI selection algorithm is designed to minimize heterogeneity, however it is not possible to completely eliminate gradients and contributions from objects that are not coplanar with the plasma membrane and focal plane. Therefore, simulations are useful for capturing the *lower limit* on the expected precision. In principle, larger ROIs can be drawn or more frames can be collected to achieve arbitrary precision as is the case for simulations. However, in practice the ROI size is limited by the inherent heterogeneity of the cellular environment and the collection time is limited by chromophore stability and cellular motion.

## Discussion

The main result of this work is the demonstration that 2c-SpCA can be used to measure heteromeric interactions between fluorescently labeled proteins in living cells. We began by implementing two techniques for measuring two-color molecular brightnesses: 2c-SpIDA and 2c-SpCA. To do this, we introduced a novel (as far we know) spectral shift filter to remove the effects of intrinsic crosstalk from our spectral detector (Figs. 2–3, S3-4). In solution, we determined that 2c-SpCA is more capable of resolving a mixture of EGFP and mApple than 2c-SpIDA, especially for small sample sizes typical of measurements in cells (Fig. S6). For cellular measurements, we extended an existing algorithm to isolate homogeneous ROIs from two-color images (Fig. 5). With our approach in place for cellular studies, we resolved constitutive heterodimers with variable FRET efficiencies labeled with EGFP and mApple (Fig. 6). Then we investigated three pairs of plasma membrane proteins. We did not observe strong interaction in a transiently interacting pair, G-DRD2/A-GNG2, in comparison to non-interacting, negative control, G-HRas/A-GNG2. However, we did observe strong interaction in a known constitutively-binding pair, G-GNB1/A-GNG2 (Fig. 7). The lack of strong interaction between G-DRD2/A-GNG2 is consistent with one hypothesis that G-protein coupled receptors must be initially activated before G-proteins bind to them (30). Thus 2c-SpCA is a new entry in the toolbox of FFS techniques that has potential to test such hypotheses in live and fixed cell studies.

### 2c-SpCA versus 2c-SpIDA

We chose 2c-SpCA for our analyses of experiments in live cells because of its superior ability to resolve multicomponent mixtures (Fig. S6), simple graphical interpretation (Fig. S7), and computational efficiency (Fig. S8). An additional advantage of 2c-SpCA is its straightforward coupling to HSP analysis. HSP analysis with temporal factorial cumulants was developed so that the heterospecies and free species have analytical relationships to the factorial cumulants (29). In principle, HSP analysis could be implemented with histogram analyses, but no such analogous analytical relationships could be derived. As consequence, the molecular brightnesses of the heterospecies and free species would not have straightforward interpretations as they do for cumulant analyses.

### SpCA and other imaging-FFS methods

In the family of FFS techniques, SpCA is most closely related to number and brightness analysis (N&B) (31) and RICS (32). Like SpCA, N&B and RICS have been extended for two-color analyses (16, 33). N&B uses many frames so that each pixel can be treated as providing a unique, temporal data set. This approach has the advantage of greater spatial resolution because particle numbers and brightnesses are reported for each pixel. SpCA combines data from ROIs in a small number of frames (Fig. 5). This approach has the possibility of greater temporal resolution for measuring fast events such as ligand-induced signal transduction. Additionally, SpCA can be used to study fixed samples because fluorescence fluctuations are generated by scanning the detection volume over particles that are stochastically distributed in space. N&B is analogous to performing a set of fixed-point measurements in parallel, therefore diffusion of particles through detection volumes corresponding to each pixel is required to measure molecular brightnesses.

RICS, like SpCA, analyzes data from regions of interest. RICS can measure dynamics (i.e. diffusivity) by considering correlations at different pixel lags. SpCA only considers cumulants with no lag. The amplitude of the autocorrelation function in RICS (and other correlation techniques) is related to the factorial cumulants by: *G*(0,0) = *κ*_[2]_/*κ*_[1]_^2^ (4, 34). Although SpCA does not provide dynamic information, it is more computationally efficient by only calculating the values necessary to find the amplitudes of the auto- and cross-correlation functions (Fig. S3A). This approach simplifies fitting and does not require a fit model of diffusion.

Regardless of the technique, it is necessary to address non-ideal detector effects to make accurate measurements in FFS. We were able to overcome intrinsic crosstalk in our multianode PMT by introducing a SSF as an additional image processing step prior to fluctuation analysis (Figs. 2–3, S3-4). The 1c-SpIDA was developed using a single analog detector (11). This required characterization of readout noise in a calibration experiment and modification of the model intensity distribution histogram to account for signal broadening. Similar calibration is performed for N&B analysis with analog detectors (35). Non-ideal detector effects in RICS (and other correlation techniques) are typically dealt with implicitly by excluding *G*(0) from the fitting routine (Fig. S3A).

### Limitations

We were able to resolve molecular interactions in living cells, however the SNR is limiting (Figs. 6–7). FFS measurements in living cells are constrained by the photophysics of the chromophores. The limiting factors on the measurements in the experiments presented here were the brightness, photostability, and maturation of mApple (Figs. 4, 6E, S5). The shortcomings of red fluorescence proteins, including mApple, in FFS are well known and have been studied extensively (36–40). More recently developed red fluorescent proteins have been developed and evaluated for use in FFS and could provide SNR improvements for multicolor FFS experiments (41–45), although a consensus choice red fluorescent protein remains elusive.

## Author Contributions

D.J.F., A.G.G., P.W.W., and D.W.P designed the research. D.J.F. and A.U. performed the research. D.J.F. and A.G.G. contributed analytic tools. D.J.F. analyzed data and wrote the manuscript.

## Acknowledgements

This work was supported by NIH grants DK08064, DK098659, and DK115972 (to D.W.P.) and a Natural Sciences and Engineering Research Council of Canada Discovery Grants RGPIN-2017-05005 (to P.W.W) and RGPIN-2019-06507 (to A.G.G.). Some experiments were supported by the Washington University Center for Cellular Imaging with the support provided by the Diabetes Research Center (DK20579) at Washington University. D.J.F. was supported by NIH grants T32EB14855 and T32DK108742.

