## Supplemental Methods for "Two-Color Spatial Cumulant Analysis Detects Heteromeric Interactions between Membrane Proteins"

**Supplementary Methods**

*2c-SpIDA.* 2c-SpIDA fits the experimental bivariate photon counting histogram, $H_{exp}(k_A, k_B)$, from imaging data with a model histogram, $H_{fit}(\bar{\varepsilon}_A, \bar{\varepsilon}_B, \bar{N}; k_A, k_B)$. $k_A$ and $k_B$ are the number of photons counted in Channels A and B, respectively. $\bar{\varepsilon}_A$ and $\bar{\varepsilon}_B$ are the molecular brightnesses in Channels A and B, respectively. We report brightness values as counts per pixel per molecule (cpm). $\bar{N}$ is the average number of fluorophores per detection volume. The lengths of $\bar{\varepsilon}_A$, $\bar{\varepsilon}_B$, and $\bar{N}$ are determined by the number of species used to calculate the model histogram. Fitting is performed by minimization of the reduced-$\chi^2$:

$$\chi^2 = \frac{\sum_{k_A,k_B}\left(M\frac{H_{exp}(k_A,k_B) - H_{fit}(\bar{\varepsilon}_A\,,\bar{\varepsilon}_B,\bar{N};k_A,k_B)}{\sigma_{k_A,k_B}}\right)^2}{\rho} \qquad (S1)$$

The $M$ is the number of pixels used to generate $H_{exp}(k_A, k_B)$. $\sigma_{k_A,k_B}$ is the estimated uncertainty of $H_{exp}(k_A, k_B)$ defined as $\sigma_{k_A,k_B} = (M \cdot H_{exp}(k_A, k_B)(1 - H_{exp}(k_A, k_B)))^{1/2}$. $\rho$ is the number of degrees of freedom.

We used the model of bivariate photon counting histograms derived by Chen et al (1). Although this model was developed for experiments where the excitation position was fixed, we were able to use it without modification under two conditions: 1) the pixel dwell time is much smaller than the characteristic diffusion time of the samples being studied, and 2) the distance swept per pixel is much smaller than the size of the point spread function (PSF). A three-dimensional gaussian (3DG) model of the PSF adequately described our data.

*2c-SpCA.* 2c-SpCA, like 2c-SpIDA, uses a model developed for describing fixed point measurements (2), but applies it to pixel values from imaging data. Bivariate factorial cumulants are related to molecular brightness and particle number density by (3):

$$\kappa_{[i,j]} = \gamma_{i+j} \cdot N \cdot {\varepsilon_A}^i \cdot {\varepsilon_B}^j \qquad (S2)$$

$\gamma_{i+j}$ are geometric factors related to the shape of the PSF. $\gamma_1$ is unity by convention. $\gamma_2$ was set to $2^{-3/2}$ for measurements in solution and 0.5 for measurements on the plasma membrane in cells. For multiple molecular species ($S$ total species), the contributions of each species ($s$) to the factorial cumulants are additive:

$$\kappa_{[i,j]} = \sum_{s=1}^{S} \gamma_{i+j} \cdot N_s \cdot {\varepsilon_{s,A}}^i \cdot {\varepsilon_{s,B}}^j \qquad (S3)$$

For our analyses, we use first and second order bivariate factorial cumulants: $\kappa_{[1,0]}$, $\kappa_{[0,1]}$, $\kappa_{[2,0]}$, $\kappa_{[1,1]}$, and $\kappa_{[0,2]}$. Again, $\bar{\varepsilon}_A$, $\bar{\varepsilon}_B$, and $\bar{N}$ are determined by $\chi^2$ minimization:

$$\chi^2 = \frac{1}{\rho}\sum_{i,j} \frac{\left(\hat{\kappa}_{[i,j]} - \kappa_{[i,j]}(\bar{\varepsilon}_A, \bar{\varepsilon}_B, \bar{N})\right)^2}{Var\left[\hat{\kappa}_{[i,j]}\right]} \tag{S4}$$

$\hat{\kappa}_{[i,j]}$ and $\kappa_{[i,j]}$ are experimental and fitted factorial cumulants, respectively. $Var[\hat{\kappa}_{[i,j]}]$ is the estimated variance of experimental factorial cumulants determined by moments-of-moments technique (2–4).

Ideal models of histograms and factorial cumulants are perturbed by non-ideal detector effects. We dealt with one effect, intrinsic crosstalk, using the spectral shift filter described below. We also measured afterpulsing in individual detector bins using methods described in (5). The afterpulsing probability across all bins was consistently near 0.001, so we used this value to make afterpulsing corrections to modeled factorial cumulants when fitting data from cellular measurements (3). In solution, molecular brightnesses were much higher due to the greater excitation intensity, therefore afterpulsing had a negligible effect and was not included in our model. Deadtime was not considered in these studies because with a multianode detector it is improbable that multiple photons will arrive in a single bin with near simultaneity.

*HSP analysis*. To analyze imaging data from cells containing multiple molecular species, we invoked heterospecies partition (HSP) analysis to reduce the number of free parameters when fitting. Our approach largely follows Wu et al (6) with the exception that we explicitly define the heterospecies brightness as having component brightnesses of single chromophore controls. We define a normalized brightness, $\bar{n}_c$, for each chromophore, $c$:

$$\bar{n}_c = (\varepsilon_{c,A}, \varepsilon_{c,B})/\varepsilon_{c,total} \tag{S5}$$

Where: $\varepsilon_{c,total} = \varepsilon_{c,A} + \varepsilon_{c,B}$. $\bar{n}_c$ contains the spectral character of $c$ independent of the molecular brightness. We use G and R to represent EGFP and mApple, respectively. We then write a chromophore specific expression of the molecular brightness for the heterospecies ($H$):

$$\bar{\varepsilon}_H = \left(\varepsilon_{H,G}, \varepsilon_{H,R}\right) \tag{S6}$$

$\varepsilon_{H,G}$ is the 'EGFP-like' brightness of the heterospecies and $\varepsilon_{H,R}$ is the 'mApple-like' brightness of the heterospecies. Conventionally, molecular brightness is written so that each component corresponds to a detection channel. We have written an alternative form where each component corresponds to a chromophore. The chromophore-specific and channel-specific forms are related by a linear transform defined by $\bar{n}_c$. The brightness of the free species ($F$) can be written as $\bar{\varepsilon}_F = \left(\varepsilon_{F,G}, 0\right)$ or $\bar{\varepsilon}_F = (0, \varepsilon_{F,R})$. The former is preferred if EGFP chromophore is more abundant and the latter is preferred if mApple is more abundant. Fitting is still performed as in Eq. 4, with the exception that $\bar{n}_G$ and $\bar{n}_R$ are additional fixed parameters used to relate the chromophore-specific brightnesses to the channel-specific factorial cumulants. In the standard formulation of HSP, only the more red-shifted chromophore can correspond to the free species. Here, either chromophore can be more abundant and therefore act as the free species. Normalized brightnesses of the heterospecies can be found by dividing by the monomeric control brightnesses:

$$b_G = \frac{\varepsilon_{H,G}}{\varepsilon_{mon,G}}\ , b_R = \frac{\varepsilon_{H,R}}{\varepsilon_{mon,R}} \qquad (S7)$$

$\varepsilon_{mon,G}$ and $\varepsilon_{mon,R}$ are the total brightnesses of EGFP and mApple monomeric controls. $b_G$ and $b_R$ are both equal to 1 for an ideal heteromer with 1:1 stoichiometry.

*Simulations.* We performed Monte Carlo simulations to generate photon counting data for fluorescent particles randomly distributed in a two-dimensional space. The total area simulated was eight times the detection volume area by the $e^{-2}$ radius of a two-dimensional Gaussian. This was to account for photons originating particles outside the detection volume. The number of photons generated for each pixel are generated as follows:

1. Determine the number of particles ($N$) in the simulated area by generating a Poissonian random number:

$$N = Pois(8 \cdot \langle N \rangle) \qquad (S8)$$

   $\langle N \rangle$ is the average number of particles per detection area for a particular molecular species.

2. The radial position ($r$) of each particle ($i$) in the 2D volume was determined by the inverse distribution method (7) so that:

$$r_i = 2\xi^{1/2} \qquad (S9)$$

   Where $\xi$ is a uniformly distributed random number between 0 and 1.
3. The *average* total number photons detected for each particle at position $r_i$ is determined by:

$$\langle n_i \rangle = \varepsilon_{tot} \cdot \exp\left(-2{r_i}^2\right) \qquad (S10)$$

   $\varepsilon_{tot}$ is the total molecular brightness across both channels.
4. The number of photons detected for each particle is determined by again generating a Poisson random number:

$$n_i = Pois(\langle n_i \rangle) \qquad (S11)$$

5. If two channels are to be simulated, $n_i$ are decomposed into two channels by generating a binomial random number:

$$n_{i,A} = Binom(n_i, p_A) \qquad (S12)$$
$$n_{i,B} = n_i - n_{i,A}$$

   $p_A$ is the probability of a detected photon originating from chromophore $i$ being detected in channel A.
6. The total number of photons detected in each channel are found by summing the contributions for each particle:

$$(n_A, n_B) = \left(\sum_i n_{i,A}\, , \sum_i n_{i,B}\right) \qquad (S13)$$

7. If multiple molecular species are simulated their contributions to the simulated photon counts are additive.

The molecular brightnesses of EGFP and mApple were set to [0.085, 0.0053] cpm and [0.00095, 0.037] cpm, respectively, to reflect measurements made in live cells. The heteromer brightness was set to [0.08595, 0.0423]. Simulations were performed so that the total number of mApple molecules would be double the total number of EGFP molecules with 0%, 50%, or 100% heteromerization:

| % Heteromerization | $\langle N_{EGFP} \rangle$ | $\langle N_{mApple} \rangle$ | $\langle N_{Het} \rangle$ |
|---|---|---|---|
| 0 | 50 | 100 | 0 |
| 50 | 25 | 75 | 25 |
| 100 | 0 | 50 | 50 |

**Table S1.** Average number of molecules per detection area for three molecular species simulated with different degrees of heteromerization.

The factorial cumulants of simulated photon count data were fit with a two-component model coupled to HSP analysis described above.
