## Supplemental figures for "Two-Color Spatial Cumulant Analysis Detects Heteromeric Interactions between Membrane Proteins"

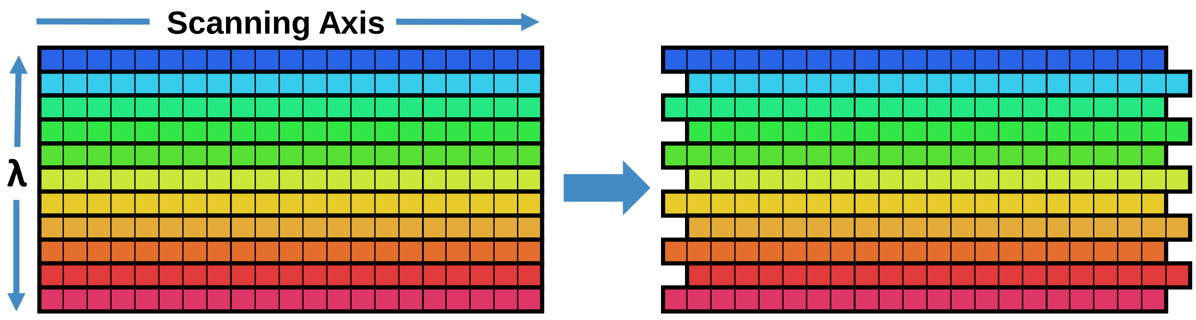


**FIGURE S1.** Schematic of spectral shift filter (SSF).

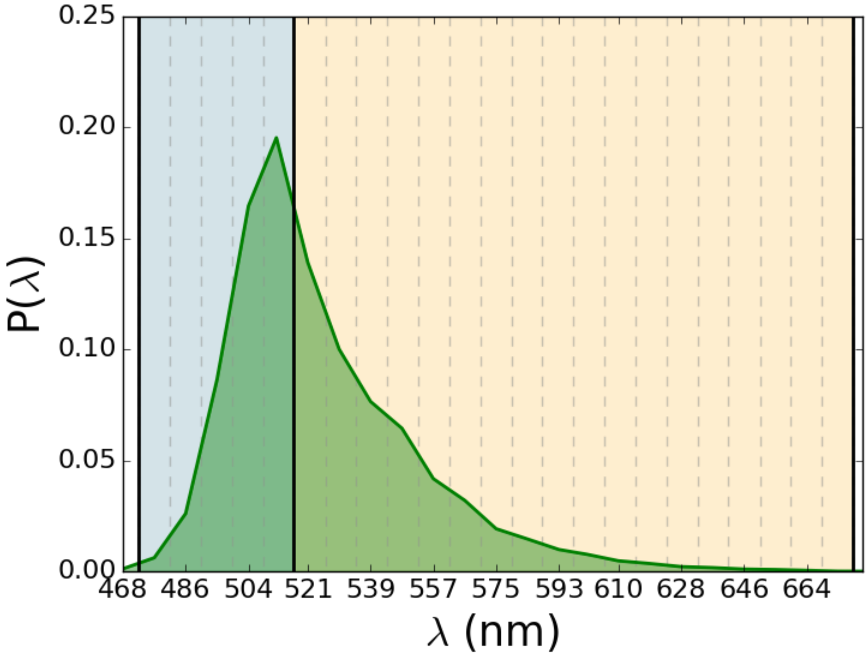


**Figure S2.** Emission spectrum of EGFP with two-channel configuration. Individual spectral bins are divided by gray dashed lines. Channels are separated by solid black lines. Channel A (472-517 nm) is highlighted in blue. Channel B (517-677 nm) is highlighted in yellow. The spectrum was normalized so that the area under the curve would be unity.

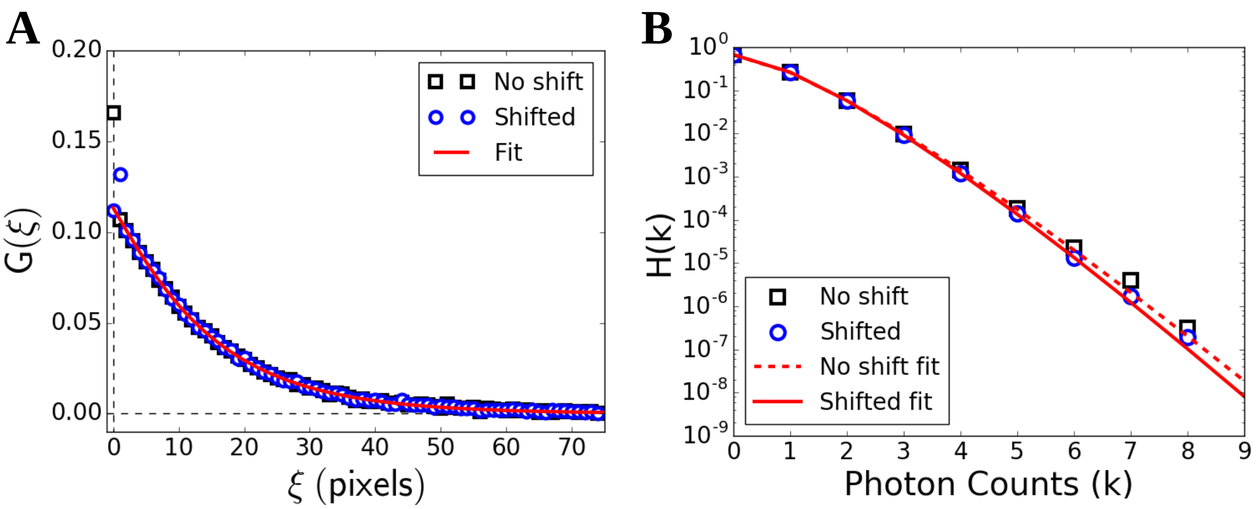


**FIGURE S3.** Effect of spectral shift filter on SACF and 1c-SpIDA for 60 frames of ~10 nM EGFP. **(A)** SACF on scanning axis (ψ=0) with no filter applied (black squares), with shift filter applied (blue circles), and fit to unfiltered data excluding G(0) (red line). **(B)** Photon counting histograms for unshifted and SSF data. Fits to unshifted and shifted histograms are shown with dashed and solid lines, respectively. Reduced-$\chi^2$ for unshifted and unshifted fits were 4.67 and 1.43, respectively.

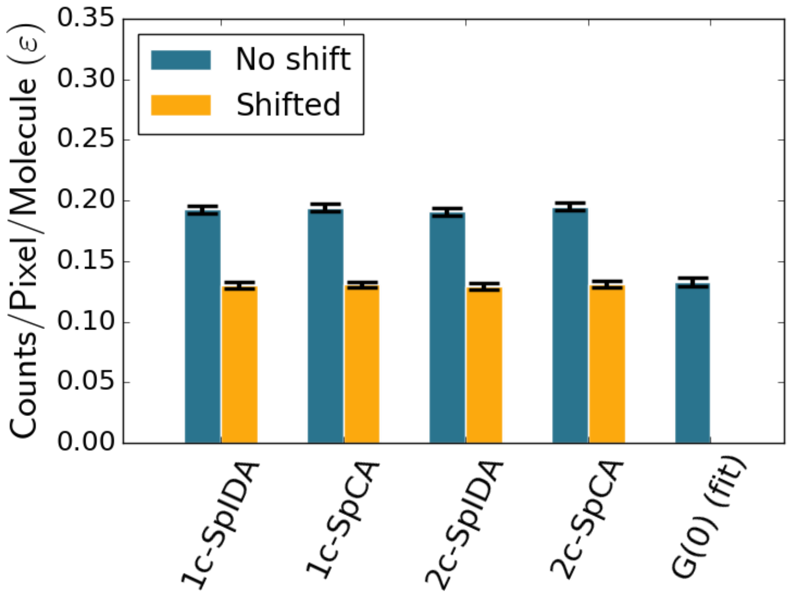


**FIGURE S4.** Brightness of ~10 nM aqueous EGFP determined by multiple methods. The error bars denote one standard deviation.

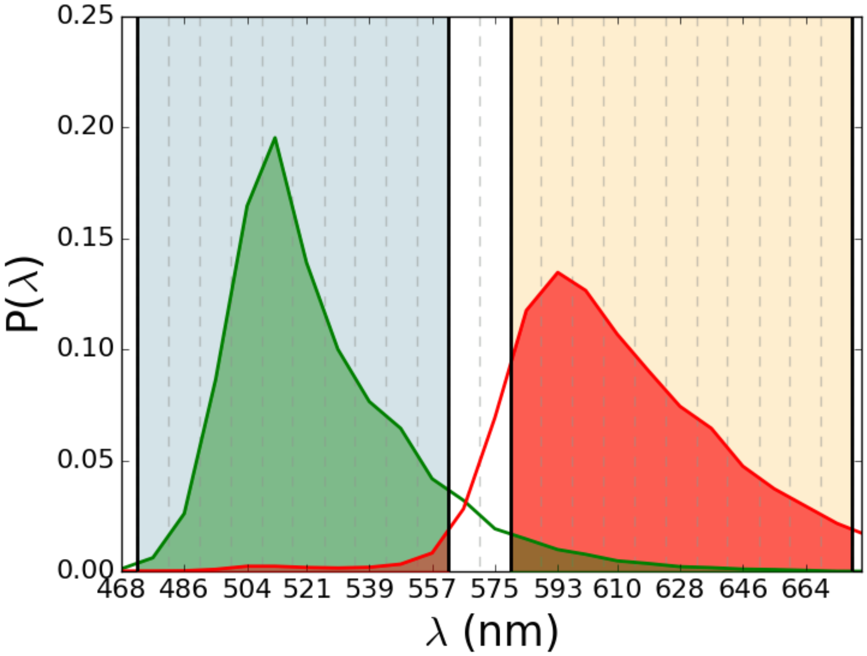


**Figure S5.** Emission spectra of EGFP and mApple with two-color channel configuration. Individual spectral bins are divided by gray dashed lines. Channels are separated by solid black lines. Channel A (472-562 nm) is highlighted in blue. Channel B (579-677 nm) is highlighted in yellow. EGFP and mApple emission spectra are highlighted with green and red, respectively. Spectra were normalized so that the areas under the curves would be unity.

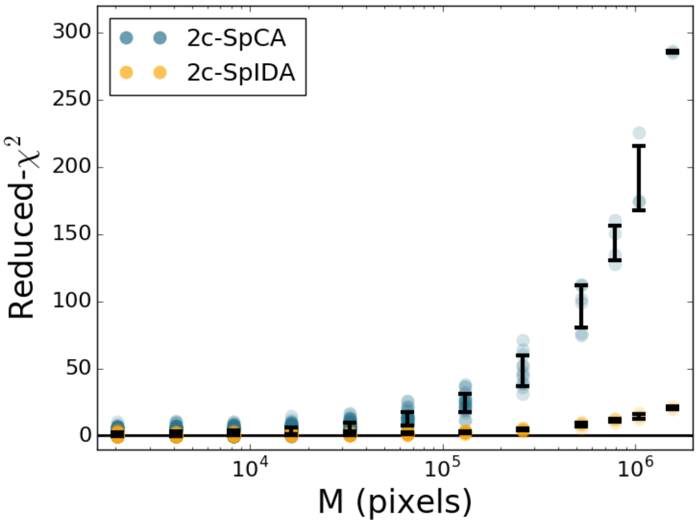


**Figure S6.** Reduced-$\chi^2$ generated by single-component fit. 60 images of ~10 nM EGFP and mApple mixture were iteratively subdivided into smaller sample sizes (M). Single-component 2c-SpCA and 2c-SpIDA were applied to each segment.

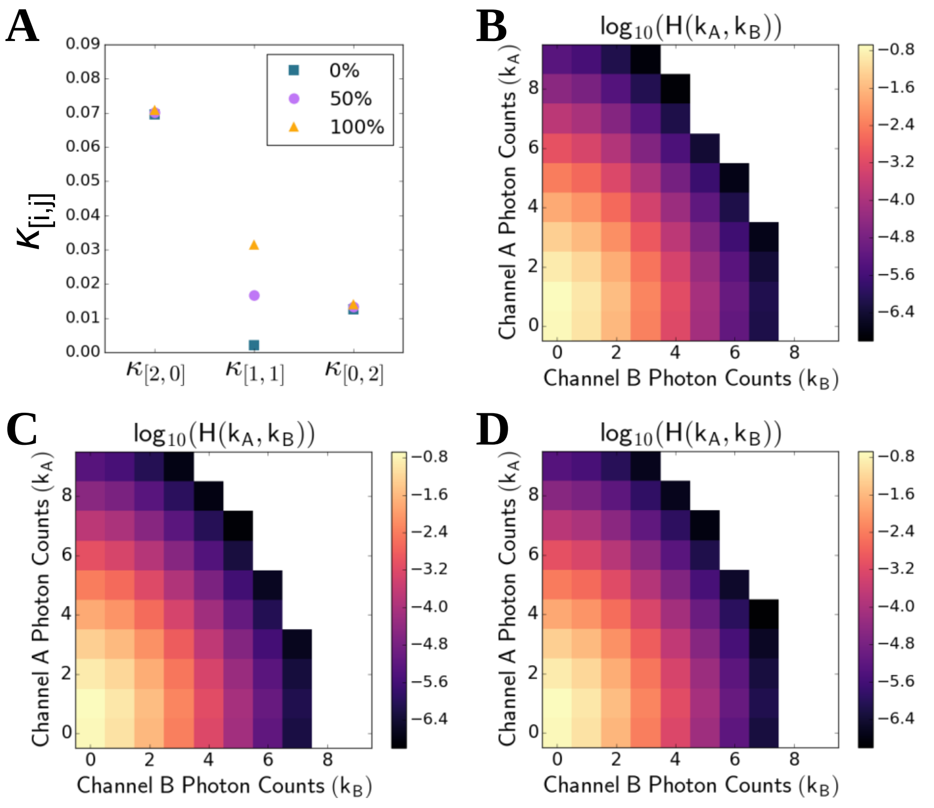


**Figure S7.** Directly calculated example factorial cumulants and intensity distributions for different degrees of heteromerization. 0% corresponds to $\overline{N}_{EGFP} = 10$, $\overline{N}_{mApple} = 10$, and $\overline{N}_{Het} = 0$. 50% corresponds to $\overline{N}_{EGFP} = 5$, $\overline{N}_{mApple} = 5$, and $\overline{N}_{Het} = 5$. 100% corresponds to $\overline{N}_{EGFP} = 0$, $\overline{N}_{mApple} = 0$, and $\overline{N}_{Het} = 10$. $\overline{N}$ are reported as average particle number per beam volume. **(A)** Second order factorial cumulants for 0%, 50%, and 100% heteromerization. **(B)-(D)** Intensity distribution for 0%, 50%, and 100% heteromerization, respectively.

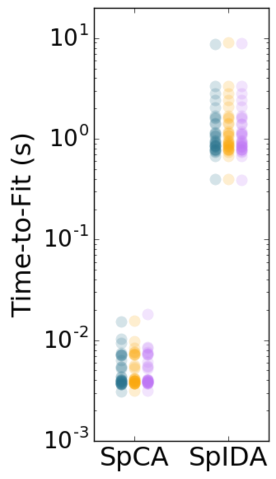


**Figure S8.** Execution time of 2c-SpCA and 2c-SpIDA fitting the same data from cells expressing EGFP-HRas. The same initial guesses were used for each fitting procedure. Three repetitions were performed, shown in blue, yellow, and pink, respectively.

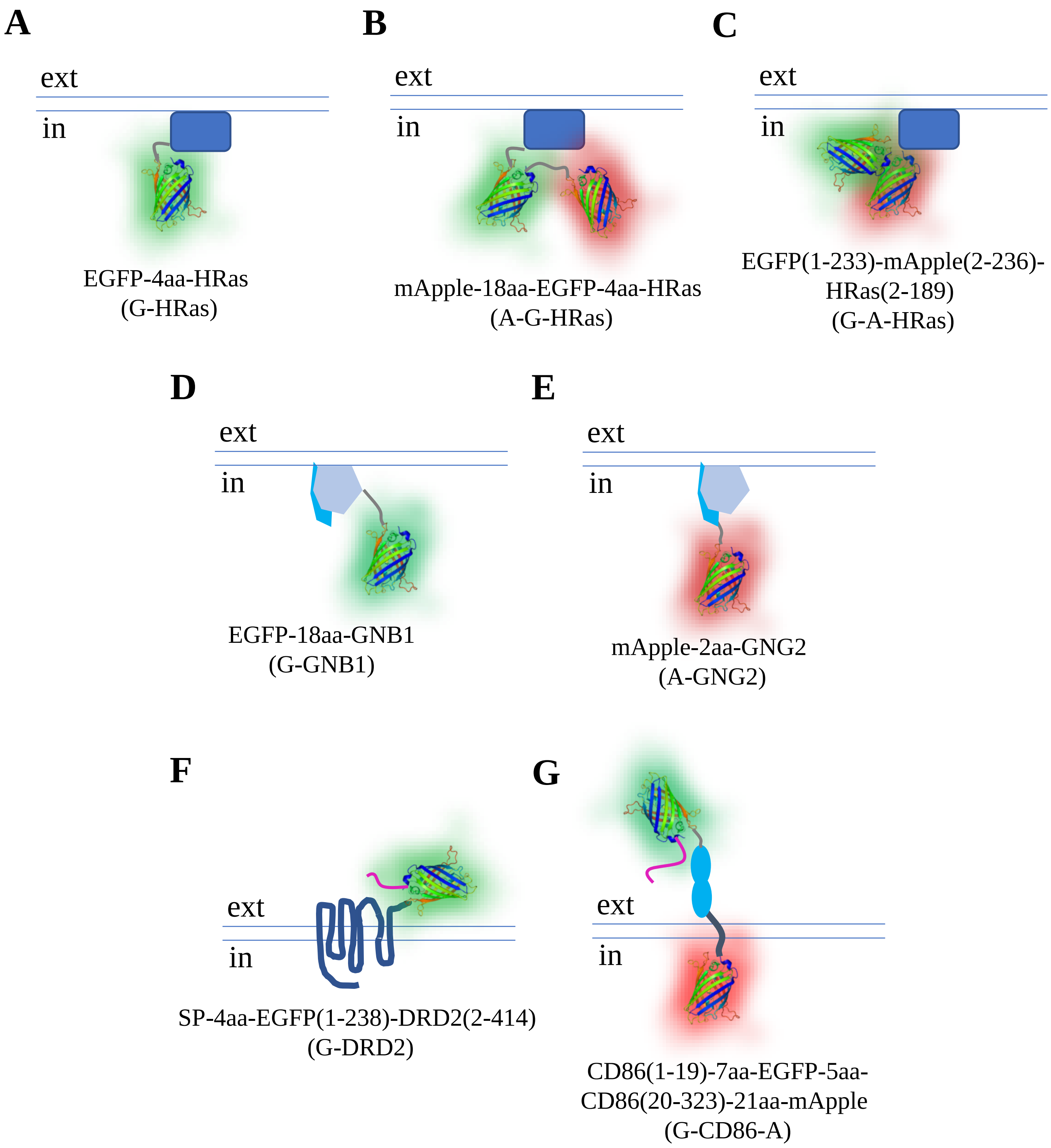


**Figure S9.** Fluorescent protein fusion constructs encoded by plasmids used in these studies. The drawings are not to scale and emphasize the distance between chormophores, presence of linker peptides, and relationship to the plasma membrane. EGFP and mApple are shown as beta-barrel ribbon structures highlighted with green and red, respectively. Linker peptides are gray. Signaling peptides are magenta. Abbreviations used in the text are shown in parentheses.

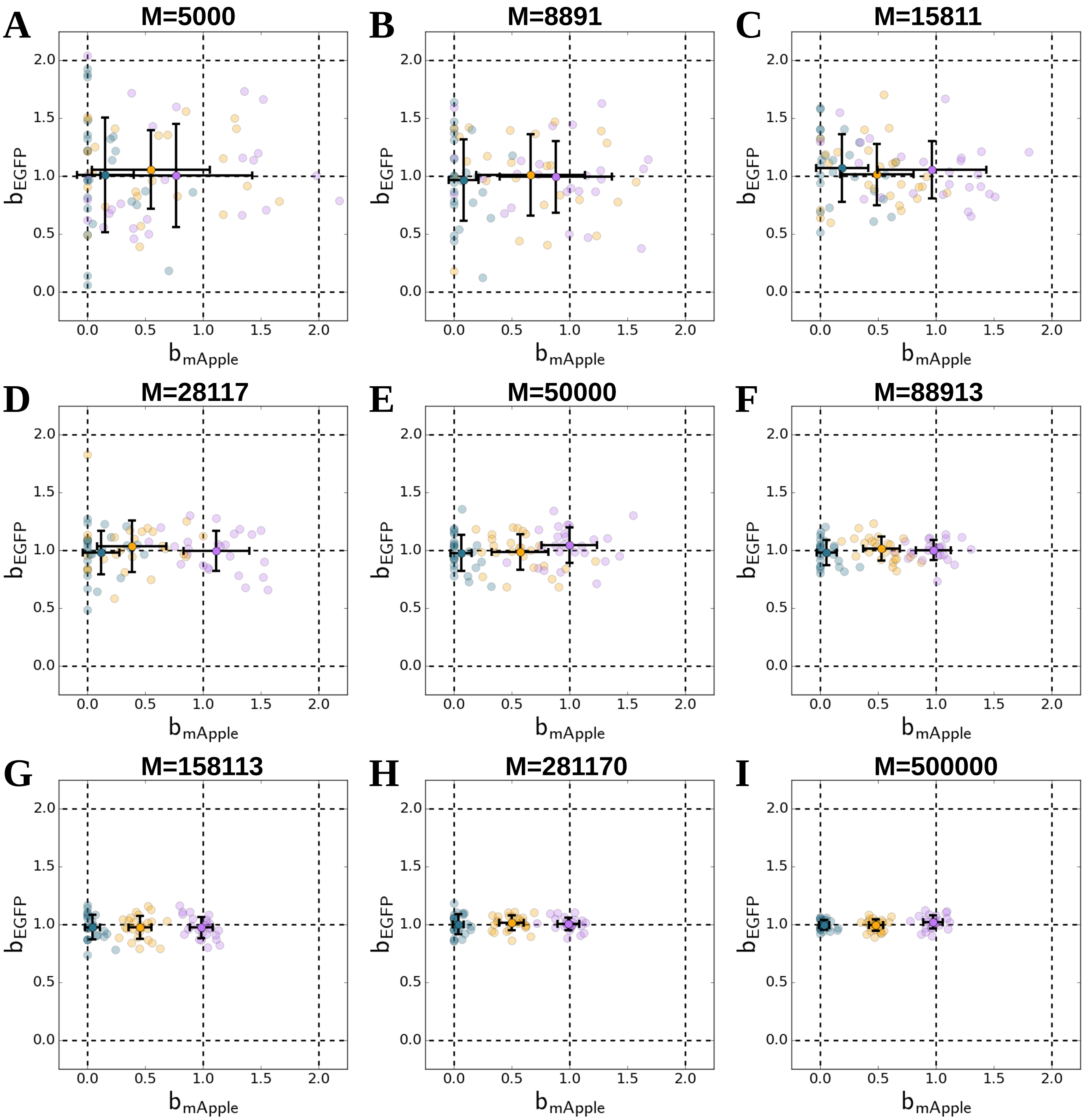


**Figure S10.** Simulations of EGFP and mApple with different degrees of heteromerization. Simulations were performed so that the total number of EGFP and mApple molecules per detection volume would be 50 and 100, respectively. Normalized brightness plots are shown for 25 simulations/parameter set. Panels **(A)-(I)** correspond to M=5000, 8891, 15811, 28117, 50000, 88913, 158113, 281170, and 500000 pixels, respectively. Blue, yellow, and pink markers correspond to 0%, 50%, and 100% heteromerization of EGFP, respectively.
